# Upscaling production of immunogenic poliovirus virus-like particles in *Pichia Pastoris* by controlled fermentation

**DOI:** 10.1101/2024.11.07.622482

**Authors:** Lee Sherry, Keith Grehan, Mohammad W Bahar, Jessica J Swanson, Helen Fox, Sue Matthews, Sarah Carlyle, Ling Qin, Claudine Porta, Steven Wilkinson, Suzanne Robb, Naomi Clark, John Liddell, Elizabeth E Fry, David I Stuart, Andrew J Macadam, David J Rowlands, Nicola J Stonehouse

## Abstract

The success of the poliovirus (PV) vaccines has enabled the near-eradication of wild PV, however, their continued use post-eradication poses a significant threat to maintaining a polio-free world. Recombinant virus-like particles (VLPs) that lack the viral genome remove this risk. Here, we demonstrate the production of PV VLPs for all three serotypes by controlled fermentation using *Pichia pastoris*. We determined the cryo-EM structure of a new PV-2 mutant, termed SC5a, in comparison to PV-2 SC6b VLPs described previously and investigated the immunogenicity of PV-2 SC5a VLPs. Finally, a trivalent immunogenicity trial using bioreactor-derived VLPs of all three serotypes in the presence of Alhydrogel adjuvant, showed that these VLPs outperform the current IPV vaccine in the standard vaccine potency assay, offering the potential for dose-sparing. Overall, these results provide further evidence that yeast-produced VLPs have the potential to be a next-generation polio vaccine in a post-eradication world.

## Introduction

Poliovirus (PV) is the causative agent of poliomyelitis, a highly infectious disease, which can cause paralysis and can be fatal. Following the launch of the Global Polio Eradication Initiative (GPEI) in 1988 there has been >99% reduction in the number of paralytic poliomyelitis cases globally, with wild-type (wt) PV-1 now only endemic in Afghanistan and Pakistan^1^. This success has resulted from the widespread and controlled use of two vaccines: live-attenuated oral PV vaccine (OPV) and inactivated PV vaccine (IPV)^2^. Both vaccines target all three serotypes of PV (PV-1, PV-2 and PV-3), and although wt PV-2 and wt PV-3 were declared eradicated in 2015 and 2019, respectively^3^, wt PV-1 and pathogenic vaccine-derived viruses remain in circulation. Despite the success of these vaccines, there are biosafety concerns associated with their continued use as we move towards a ‘polio-free’ world, since both vaccines depend on the culture of infectious virus and have the potential to re-introduce PV into the environment.

IPV provides excellent protection against disease: however, like all inactivated vaccines, it is ineffective in inducing mucosal immunity. Consequently, it cannot prevent transmission of PV and can result in ‘silent’ spread within a population^4^. Furthermore, the cultivation of large amounts of infectious virus, required for the manufacture of conventional IPV, presents an inherent potential biosafety risk^5,6^. OPV has been of critical importance in the near-eradication of PV, but the attenuated virus can readily revert to virulence. This can result in vaccine-associated paralytic poliomyelitis (VAPP) which, in low vaccine coverage areas, can lead to circulating vaccine-derived PV (cVDPV)^7^. Currently, cVDPV cases outnumber wt PV cases worldwide, as recently exemplified by the detection of PV-2 VDPV sequences in environmental samples in London, UK and the identification of a paralytic polio case in the USA^1,8,9^. Furthermore, OPV can recombine with other PVs or polio-like enteroviruses during co-infection to generate novel neurovirulent chimeric viruses^10,11^. This, together with the risk of further reintroduction of infectious PV into the environment via the chronic shedding of VDPV by immunocompromised individuals, emphasises the risks concomitant with continued OPV usage as we strive towards eradication^11,12^.

PV belongs to the picornavirus species *Enterovirus C* with a 7.5 kb positive-strand RNA genome which comprises two overlapping open-reading frames (ORFs), the uORF and the major polyprotein ORF (ppORF)^13^. The ppORF is translated as a single polypeptide, containing 3 distinct regions, P1 (viral structural proteins), P2 and P3 (non-structural proteins required for proteolytic cleavage and viral replication). The P1 region is separated from the polyprotein by the self-cleaving 2A^pro^; P2 and P3 are cleaved into the mature viral proteins by viral proteases 3C^pro^ and 3CD^14^. The viral protease precursor protein, 3CD, is primarily responsible for the cleavage of P1 into the individual capsid proteins, VP0, VP3 and VP1^15,16^. During virion maturation, VP0 is cleaved into VP4 and VP2, concomitant with encapsidation of viral RNA, and results in increased particle stability^17,18^.

Mature PV virions are ∼30 nm diameter icosahedral capsids comprising 60 copies of VP1-VP4^19^. Capsid stability is enhanced by the incorporation of a host-derived lipid molecule, known as pocket factor, into a cavity within the VP1 protein^20^. Empty capsids (ECs), which are also produced during infection, are antigenically indistinguishable from mature viral particles but do not contain viral genome and do not cleave VP0^21^. ECs have potential as virus-like particle (VLP) vaccines to replace the current PV vaccines; however, recombinantly expressed wt ECs are inherently unstable and readily convert to an expanded form^21,22^. This expansion converts ECs from the native D antigenic form (D Ag) to the non-native form (C Ag)^23^. C Ag particles do not induce a protective immune response and recombinant VLP vaccines must therefore retain the D Ag conformation^22,24,25^.

It has been demonstrated by our previous work that the native conformation can be stabilised by the introduction of mutations within the viral capsid^26^. VLPs are attractive as recombinant vaccine candidates, with excellent biosafety profiles, as highlighted by the success of the licensed hepatitis B virus (HBV) and human papillomavirus (HPV) vaccines produced in yeast or insect cells^27–29^.

Recombinant PV VLPs have been produced in various expression systems, including mammalian, plant, and insect cells^26,30–35^. Recently, we have demonstrated the production of antigenically stable PV VLPs using the yeast, *Pichia pastoris*^36^. Yeast has several potential advantages as a PV VLP production system as it is highly scalable and has possibilities for technology transfer to LMICs possessing the infrastructure used to produce the HBV and HPV vaccines^37^. Our WHO-funded consortium has recently demonstrated that recombinant PV VLPs from several different expression systems, including yeast-derived VLPs, are as immunogenic as the current IPV in the rat assay used to certify vaccine batch release^38^.

Here, we demonstrate the scalability of *Pichia pastoris* as a production system to produce stabilised VLPs for all three PV serotypes using controlled fermentation. The effects of different media types and methods of induction on D Ag production for all three serotypes were investigated using small 250 mL bioreactors. The best performing conditions were then applied to 10 L bioreactor vessels and D Ag yield compared to flask-based expression. A modified PV-2 stabilised mutant, SC5a, resulted in improved D Ag yield with greater thermostability than the current IPV vaccine. We determined the cryo-EM structures of the yeast-derived PV-2 SC6b and PV-2 SC5a VLPs and investigated the immunogenicity of the PV-2 SC5a alone or with aluminium hydroxide (Al(OH)_3_) adjuvant in a rat model. In a trivalent immunogenicity trial using bioreactor-derived VLPs of all three serotypes, in the presence of adjuvant, the recombinant VLPs outperformed the current IPV vaccine in the rat model, offering the potential for dose-sparing. Overall, these results provide further evidence that yeast-produced VLPs have the potential to be next-generation polio vaccines in a post-eradication world.

## Results

### Bioreactor production of PV VLPs

We have previously reported the production and immunogenicity of yeast-derived PV VLPs using laboratory-scale shaker flask expression^36,38,39^. However, to become successful vaccine candidates it will be necessary to produce the particles using methods suitable for industrial manufacture. To this end, we explored the potential for scale-up of production using the controlled fermentation conditions achievable in bioreactors as applied to the commercial production of current HBV and HPV vaccines^29^.

Initially, we expressed previously characterised stabilised mutants for each PV serotype (termed PV-1 SC6b, PV-2 SC6b, and PV-3 SC8, respectively^26^ at small scale (∼170 mL) in an Ambr250 Bioreactor comparing temperature, media type and feed conditions (Table 1). To assess antigen production in each condition in comparison to flask-based expression, cell pellets were collected 48 h post-induction and frozen at −20℃ before processing. These were resuspended prior to homogenisation at ∼275 MPa and the resultant lysates purified through chemical precipitation and differential centrifugation steps culminating in 15 to 45% sucrose gradients (Fig. 1). VLPs were assessed in terms of antigen content using specific monoclonal antibodies, as described in Methods. To investigate PV VLP expression from *P. pastoris* under controlled fermentation conditions we trialled PV-1 SC6b using YPD/M, a low-cost simple expression media, or BMGY/BMMY, (a more complex media optimised for the extracellular secretion of recombinant proteins) under fermentation conditions at 24℃. In addition, we compared a continuous methanol feed (1.8 mL/h/L) with a bolus feed (0.1% v/v) every 24 hours (h) (Table 1). Under continuous feed conditions both the quantity and quality (D:C ratio) of VLPs produced in YPD/M were better than in BMGY/BMMY (1491 (+/- 21) D Ag/100 mL vs 876 (+/- 120) D Ag/100 mL). In bolus-fed conditions VLP yields and D:C ratios were similar in both media but the amount of D Ag/100 mL was less in comparison to the continuous-feed YPD/M condition (1491 (+/- 21) vs 877 (+/- 108) D Ag/100 mL). Thus, continuous-feed expression produced greater yields of PV VLP when cultured in YPD/M medium.

**Figure 1:**
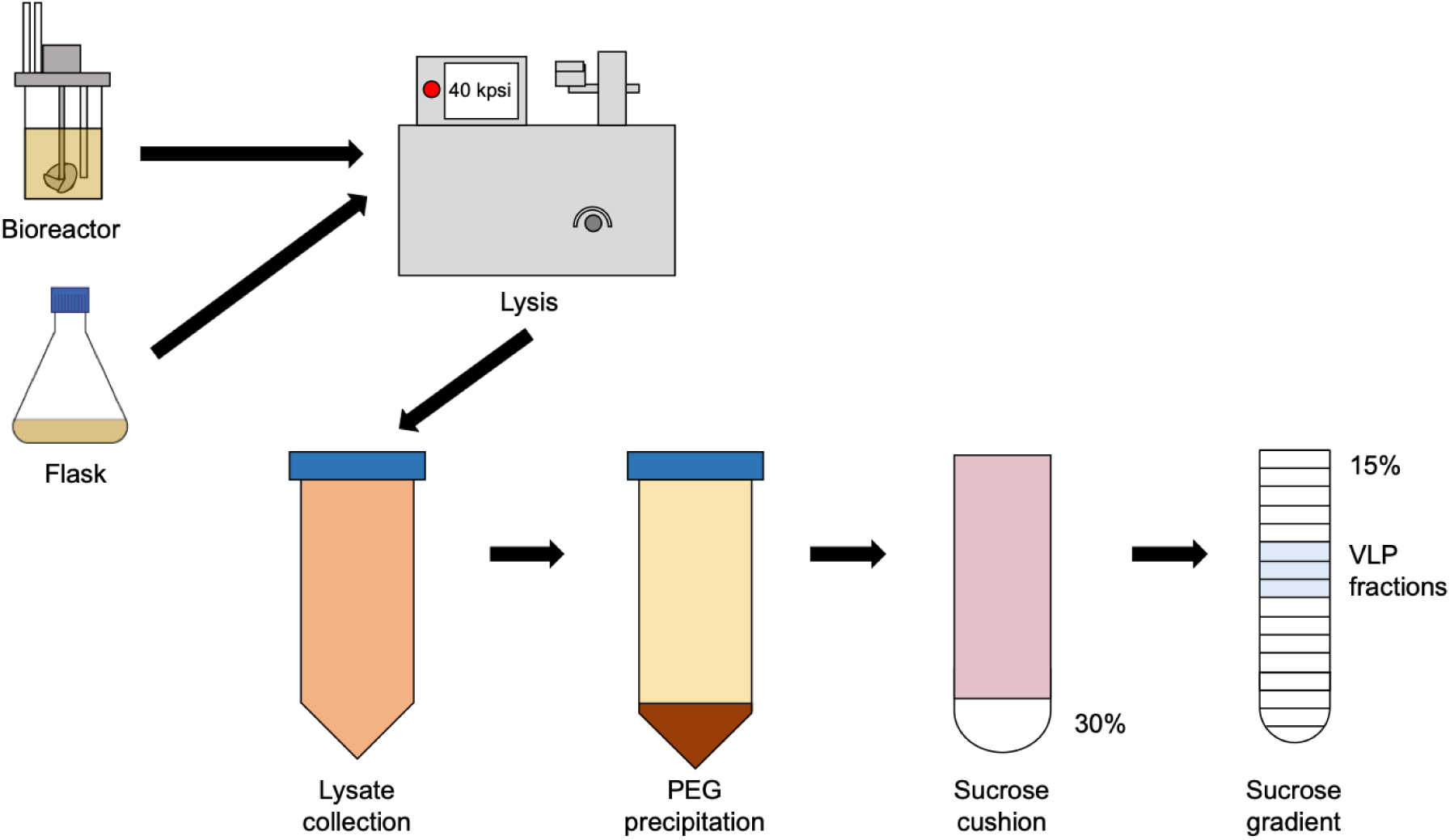
VLP purification schematic: To compare yields of yeast-derived PV VLPs produced by either flask or bioreactor mediated expression, yeast pellets are processed in the same way. Pellets derived from equivalent culture volumes were mechanically disrupted at 40 kpsi (∼275 MPa) and the cell lysates collected for further processing. Following removal of insoluble material, samples were treated with nuclease prior to a chemical precipitation step using PEG. The pelleted material was then resuspended and pelleted through 30% sucrose. The resulting pellet was resuspended in a small volume prior to sucrose density gradient ultracentrifugation. The resultant fractions were analysed for the presence of PV VLPs by ELISA.

**Table 1.** Ambr250 bioreactor conditions evaluated for each PV serotype.

| Serotype | Media | Temp (°C) | Feed Conditions | VLP Yield<br>(D Ag/100 mL<br>culture) | D:C Ag<br>Ratio |
| --- | --- | --- | --- | --- | --- |
| PV1 SC6b | YPD/M | 24 | Continuous | 1491 (+/- 21) | D >> C |
|  |  |  | Bolus | 877 (+/- 108) | D > C |
|  | BMGY/BMMY | 24 | Continuous | 876 (+/- 120) | D >> C |
|  |  |  | Bolus | 843 | D > C |
| PV2 SC6b | YPD/M | 26 | Continuous | 658 (+/- 36) | N/A |
|  |  | 28 | Continuous | 652 (+/- 40) | N/A |
|  |  | 30 | Continuous | 677 (+/- 38) | N/A |
|  | Minimal | 26 | Continuous | 23 (+/- 21) | N/A |
|  |  | 28 | Continuous | 5 (+/- 5) | N/A |
|  |  | 30 | Continuous | 2 (+/- 1) | N/A |
| PV3 SC8 | YPD/M | 26 | Continuous | 1400 (+/- 160) | D > C |
|  |  | 28 | Continuous | 1921 (+/- 4) | D > C |
|  |  | 30 | Continuous | 2677 (+/- 167) | D ≈ C |
|  | Minimal | 26 | Continuous | 7 (+/- 2) | D = C |
|  |  | 28 | Continuous | 32 (+/- 28) | D > C |
|  |  | 30 | Continuous | 4 | D = C |

To further optimise the yield of D Ag VLPs, both PV-2 SC6b and PV-3 SC8 constructs were expressed under continuous-feed conditions at three temperatures (26℃, 28℃ or 30℃) using two different media, YPD/M or minimal medium, which is optimised for the expression of difficult to produce proteins. However, in minimal medium D Ag yields were very low for both constructs at all temperatures and were not viable for bioreactor production of PV VLPs (Table 1). There were no significant differences in the D Ag yield for PV-2 SC6b at different temperatures in the YPD/M expression conditions. In contrast, PV-3 SC8 VLP D Ag yield increased by ∼500 D Ag/100 mL (equivalent to ∼18 human vaccine doses) when the expression temperature was increased to 28℃, whilst maintaining a similar D:C ratio. Although the D Ag/100 mL yield increased to 2677 (+/- 167) at 30℃, the D:C ratio was considerably reduced (Table 1). Therefore, in order to maximise the production of VLPs in the native conformation (D Ag) whilst keeping a favourable D:C Ag ratio, we selected continuous-feed, YPD/M at 28℃ in 10 L cultures for each PV serotype.

Following fermentation, cell pellets were processed as described in Fig. 1 for each serotype. The antigenic content was assessed by ELISA using a standard protocol established by the National Institute for Biological Standards and Control (NIBSC – now MHRA) with the current inactivated vaccine standard (termed BRP) as a positive control. As expected, the gradient profile for each serotype showed a strong D Ag peak centred on fractions 6 and 7, with levels above or equal to the BRP standard (Fig 2A). Peak fractions from these gradients were concentrated using 100 kDa centrifugal concentrators and assessed by TEM for particle morphology. Near-uniform particles of ∼30 nm diameter were seen for each serotype, consistent with previous EM images of PV virions and empty capsids and yeast-derived PV-1 VLPs^40^ (Fig. 2B).

**Figure 2:**
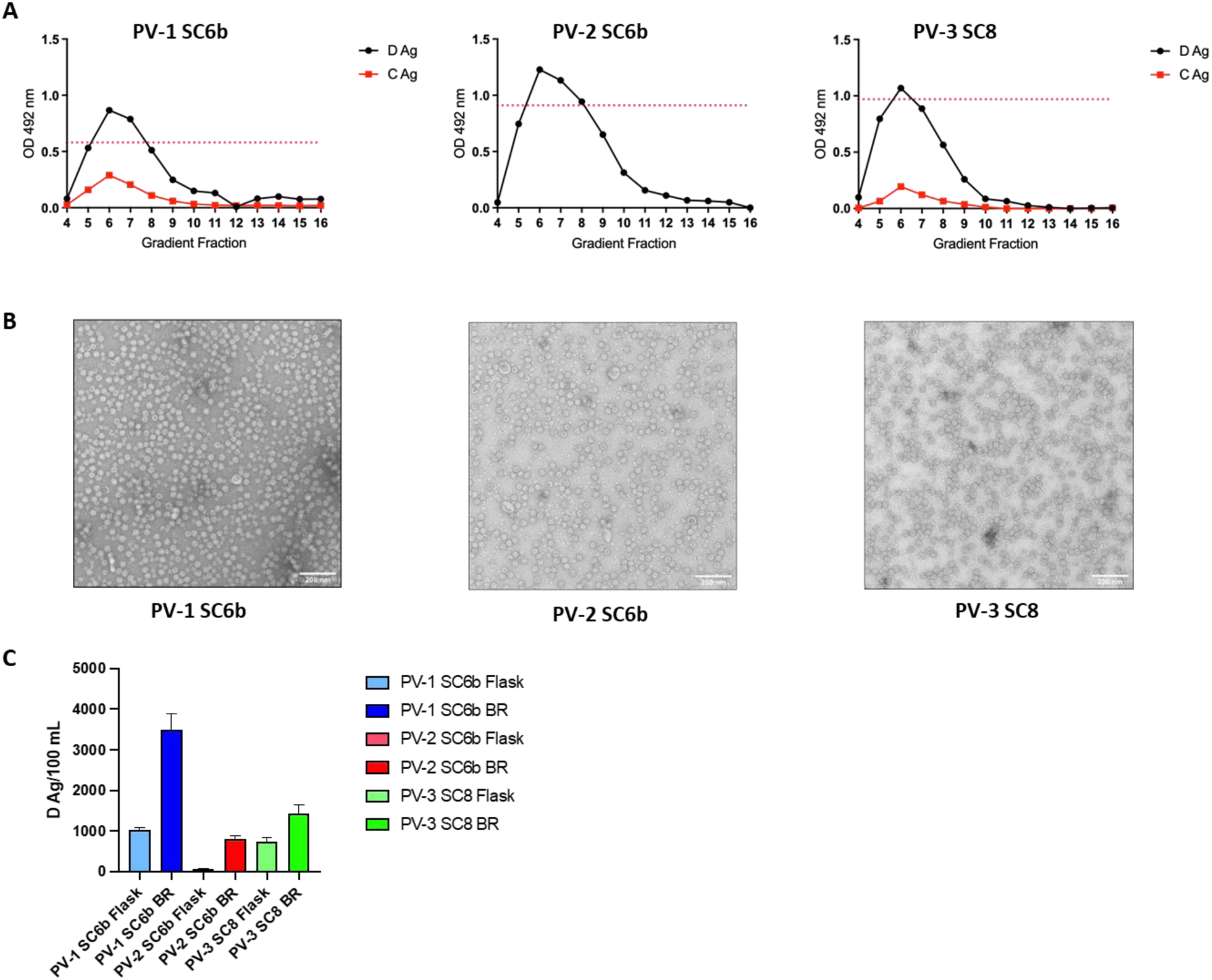
Production of PV VLPs under fermentation conditions: **A**. Antigenicity of PV VLPs. Reactivity of gradient fractions using PV serotype-specific monoclonal antibodies, for D antigen (MAb 234, 1050, and 520 for PV-1, PV-2 and PV-3, respectively) or C antigen (MAb 1588 and 517.3 for PV-1 and PV-3, respectively) in ELISA. The pink dashed line represents the positive control, BRP, for the D antigen ELISA. OD at λ=492 nm is represented in arbitrary units. The figure is a representative example of three separate experiments for each construct. **B**. Representative micrographs of purified PV VLPs (Scale bar shows 200 nm). **C**. Comparative analysis of VLP production in flask and bioreactor. Amount of purified D Ag produced per 100 mL culture as determined by ELISA detect using MAb 234, 1050, and 520 for PV-1, PV-2 and PV-3, respectively. OD at λ=492 nm is represented in arbitrary units (n = 3)

We compared the D Ag yield from both flask-based and bioreactor-based expression from multiple purifications (n = 3) (Fig. 2C). Significant improvements in yield were observed for each serotype when produced under controlled fermentation conditions. The yield of PV-1 SC6b D Ag was increased by ∼3 fold and that of PV-3 SC8 was improved ∼2-fold whereas the yield of PV-2 SC6b was ∼10-fold greater in comparison to flask-based expression. Overall, these results demonstrate the suitability of *Pichia pastoris* as a scalable expression system for the production of PV VLPs as future vaccine candidates.

### The PV-2 stabilised mutant SC5a improves VLP yield in both flask and bioreactor production

As yields of PV-2 SC6b were lower than those of PV-1 SC6b or PV-3 SC8 (Fig. 2C) we investigated shaker flask expression of an alternative stabilised PV-2, termed PV-2 SC5a^26,41^. ELISA of sucrose gradient fractions showed that the yield of D Ag was significantly higher for PV-2 SC5a compared to PV-2 SC6b (Fig. 3A). A peak fraction from each gradient was concentrated using 100 kDa centrifugal concentrators and examined by TEM, which showed that like PV2-SC6b, mutant PV-2 SC5a produced ∼30 nm particles as expected (Fig. 3B).

**Figure 3:**
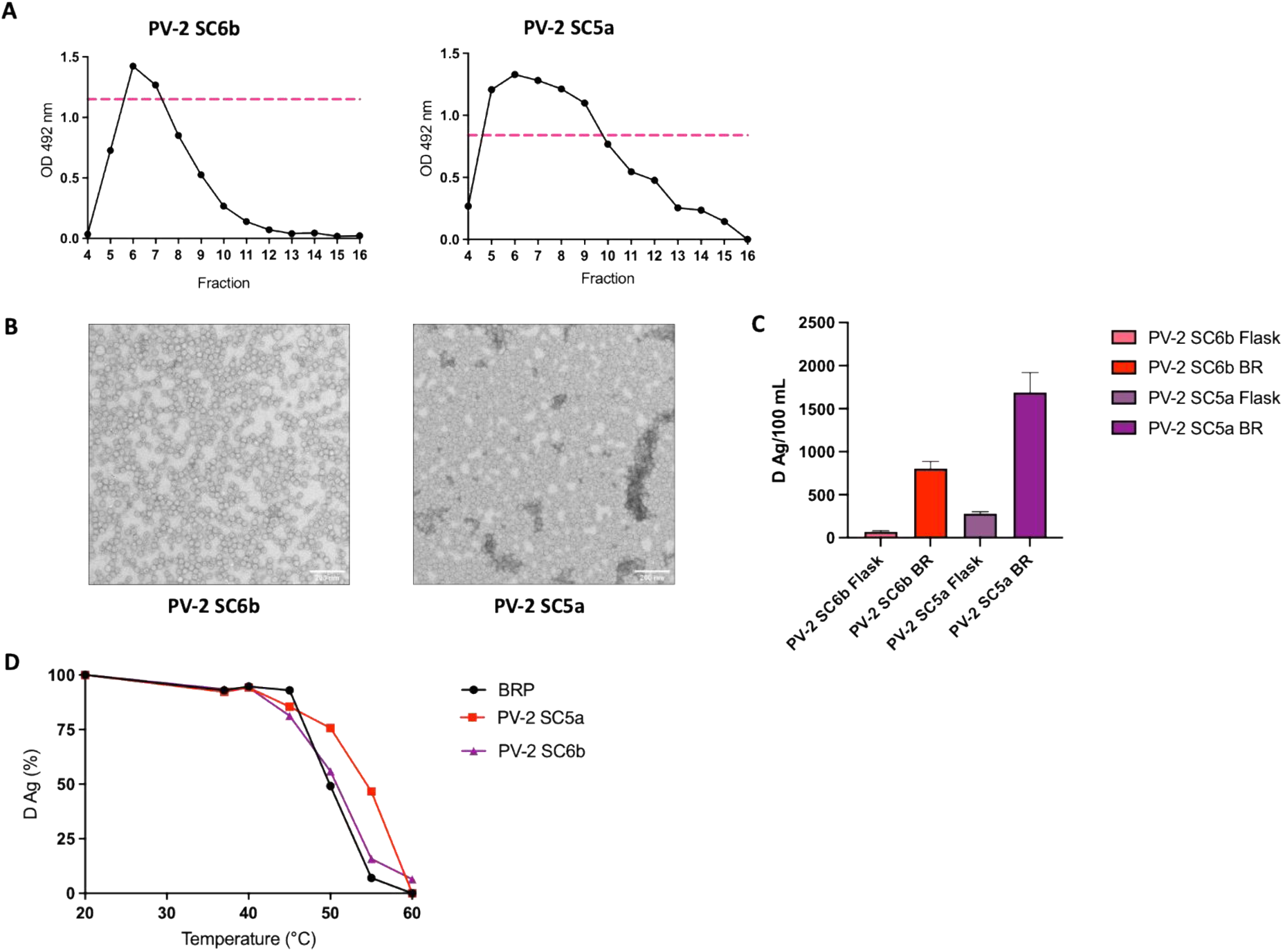
Comparison of PV-2 SC6b and PV-2 SC5a VLPs: A. Antigenicity of thermally stabilised PV-2 VLPs. Reactivity of gradient fractions using a PV-2 specific monoclonal antibody for D antigen (MAb 1050) in ELISA. The pink dashed line represents the positive control, BRP. OD at λ=492 nm is represented in arbitrary units. The figure is a representative example of three separate experiments for each variant. B. Representative micrographs of purified PV-2 VLPs (Scale bar shows 200 nm). C. Comparative analysis of PV2 VLPs production in flask and bioreactor. Amount of purified D Ag produced per 100 mL culture as determined by ELISA using MAb 1050. OD at λ=492 nm is represented in arbitrary units (n = 3) D. Reactivity of purified PV-2 SC6b and PV-2 SC5a VLPs and BRP aliquots to D-antigen specific MAb 1050 in ELISA after incubation at different temperatures for 10 minutes, normalised to corresponding aliquot incubated at 4 °C. The figure is a representative example of two separate experiments for each variant

PV-2 SC5a produced more D Ag/100 mL than PV-2 SC6b in both flask and bioreactor expression systems (3-fold and ∼2-fold respectively) (Fig. 3C). We also compared the thermal stability of PV-2 SC5a with those of PV-2 SC6b and the current inactivated vaccine, BRP, as a positive control. PV-2 SC5a was highly stable, with ∼75% D Ag remaining at 50℃ and ∼40% D Ag at 55℃, outperforming both PV-2 SC6b and BRP (Fig. 3D).

### Cryo-EM structure determination and comparison of PV-2 SC6b and PV-2 SC5a VLPs

Detailed structural analyses and comparison of the PV-2 SC6b and PV-2 SC5a VLPs from bioreactor production were performed by single particle cryogenic electron microscopy (cryo-EM). Sucrose density gradient fractions for each PV-2 VLP were pooled, concentrated and applied to EM grids that were rapidly vitrified. Data were collected as described in Methods. Data processing yielded a final set of 113,853 particles for PV-2 SC6b and 147,409 particles for PV-2 SC5a, and icosahedral reconstructions of 2.4 Å and 2.1 Å resolution, respectively (Fig. 4A, 4B and Table S1).

**Figure 4:**
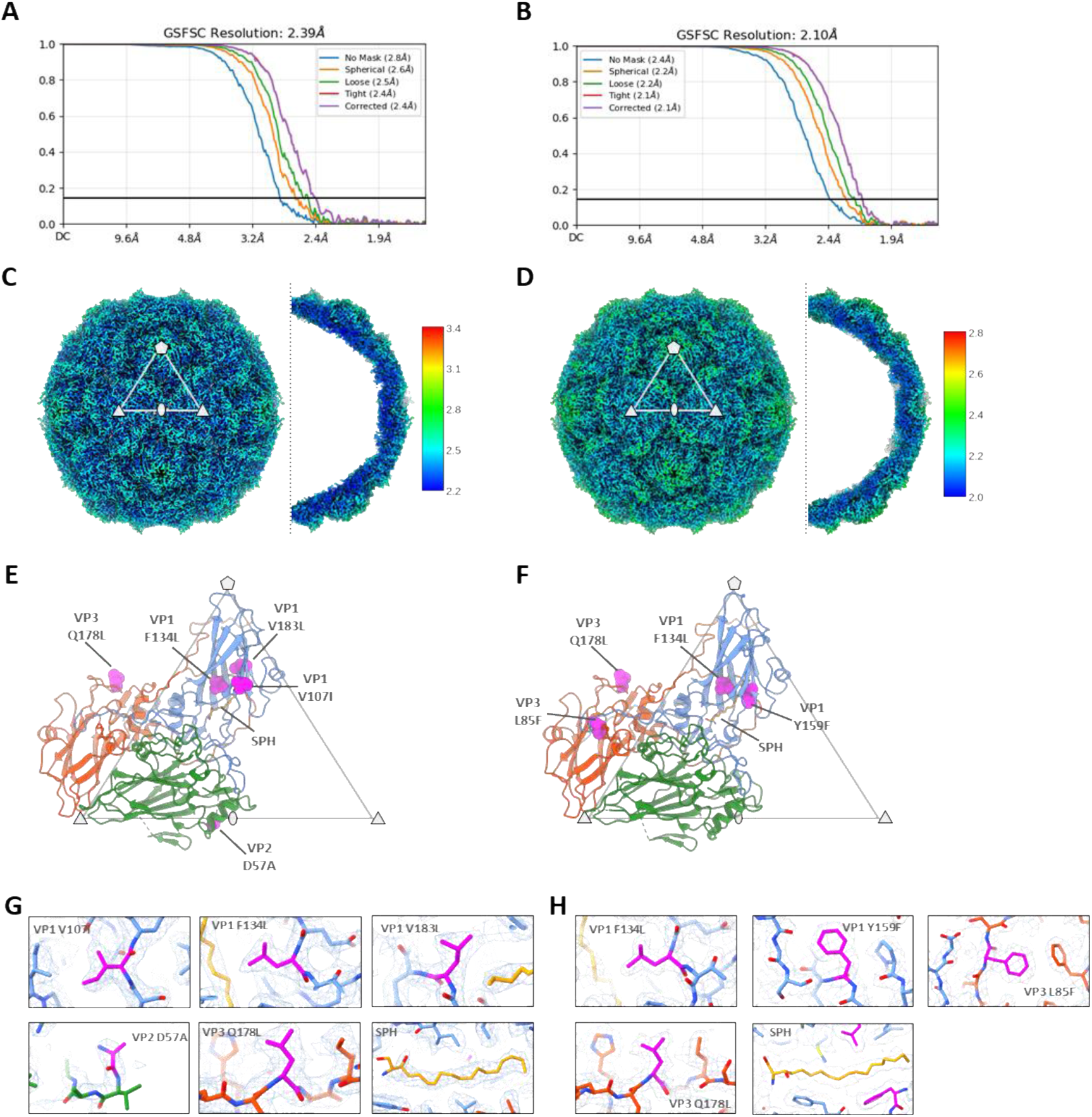
Cryo-EM analysis of bioreactor produced PV-2 SC6b and PV-2 SC5a VLPs. Gold standard Fourier shell correlation (GSFSC) curves calculated between two independent half sets of data as a function of resolution are shown for PV-2 SC6b (A) and PV-2 SC5a (B), generated after local resolution analysis in CryoSPARC. Isosurface representations of the electron potential maps for PV-2 SC6b (C) and PV-2 SC5a (D) are shown at a threshold of 4σ (σ is the standard deviation of the map). Representative five-fold, three-fold and two-fold symmetry axes are indicated with symbols, and the white triangle delimits an icosahedral asymmetric unit (AU). A central slice through half of each VLP is viewed along the icosahedral two-fold axis. Maps are coloured by local resolution (in Å) according to the colour key. Molecular cartoon representations of a single capsid protomer for PV-2 SC6b (E) and PV-2 SC5a (F) is shown within a triangle depicting an icosahedral AU, with symmetry axes labelled with symbols as in (C) and (D). Individual subunits of the capsid protomer are coloured blue (VP1), green (VP0) and red (VP3). Stabilising mutations engineered into each VLP are shown as spheres coloured magenta and labelled. For VP2 D57A sequence numbering for the mature VP2 peptide is used (equivalent to D126A in VP0 numbering). The sphingosine lipid moiety modelled into the VP1 hydrophobic pocket is shown as sticks coloured orange and labelled (SPH). Close up views of the stabilising mutations and SPH in PV-2 SC6b (G) and PV-2 SC5a (H) are depicted as sticks fitted in the cryo-EM electron potential maps, shown as a wire mesh (threshold 2σ). SPH for PV-2 SC5a is shown fitted into the map at a threshold of 1.5 σ. For PV-2 SC6b the mutation in subunit VP4 I57V (I57V in VP0) and for PV-2 SC5a the mutation in subunit VP1 T41I, were in disordered regions of the map and were not modelled.

The structures confirm that both PV-2 VLPs adopt a homogeneous D Ag conformation, with negligible or no particles observed in C Ag form (Fig. 4C, 4D). The quality of the cryo-EM electron potential maps is sufficient to resolve the stabilising mutations and confirm that they introduce no significant perturbations to the structure and do not impinge on the antigenic surface of the particle (Fig. 4E-H). Furthermore, the structure of the PV-2 SC6b VLP from yeast bioreactor material is identical in overall features and conformation to the same VLP produced in mammalian and insect cells^38^ with an rmsd in Cα atoms of 0.31 Å and 0.49 Å respectively between capsid protomers demonstrating that the fine structure is also essentially indistinguishable. As expected, the PV-2 SC5a variant also matches the PV-2 SC6b mutant closely with no major changes in conformation or antigenic sites (0.45 Å rmsd in Cα atoms between protomers) (Fig. 4E, 4F).

The observed pocket factor density in the VP1 subunit of the capsid protomer is the same in both PV-2 SC6b and PV-2 SC5a variants from bioreactor production, and closely matches that observed in PV-2 SC6b from mammalian and insect expression^38^ (Fig. 4G, 4H). Based on the extent of the observed electron potential map in the VP1 pockets of both PV-2 SC6b and PV-2 SC5a, a sphingosine lipid moiety of carbon length 18 was modelled into the maps (Fig. 4G, 4H).

In PV-2 SC6b an apparent disulphide bond is observed between residues Cys130 and Cys326 of the VP0 subunit (equivalent to Cys61 and Cys257, respectively, of the mature VP2 peptide) (Fig. S1). These cysteine residues are conserved across all serotypes of PV; however, the disulphide is not observed in either PV-2 SC5a reported in this study (Fig. 4), nor in the previously determined structures of PV-2 SC6b from mammalian and insect cell expression^38^.

### Bioreactor-derived PV-2 SC5a VLPs are immunogenic in a rat model

Following the antigenic, thermal stability and structural characterisation of the PV-2 SC5a VLPs, we assessed their immunogenicity using the pharmacopeial IPV lot release assay in Wistar rats^42^. Rats were immunised with either IPV or PV-2 SC5a with or without adjuvant (Alhydrogel 2%) at doses ranging from 1 human dose (8 D AgU) to 0.125 human dose (1 D AgU) and the resulting sera assessed for neutralising antibody titres (Fig. 5).

**Figure 5:**
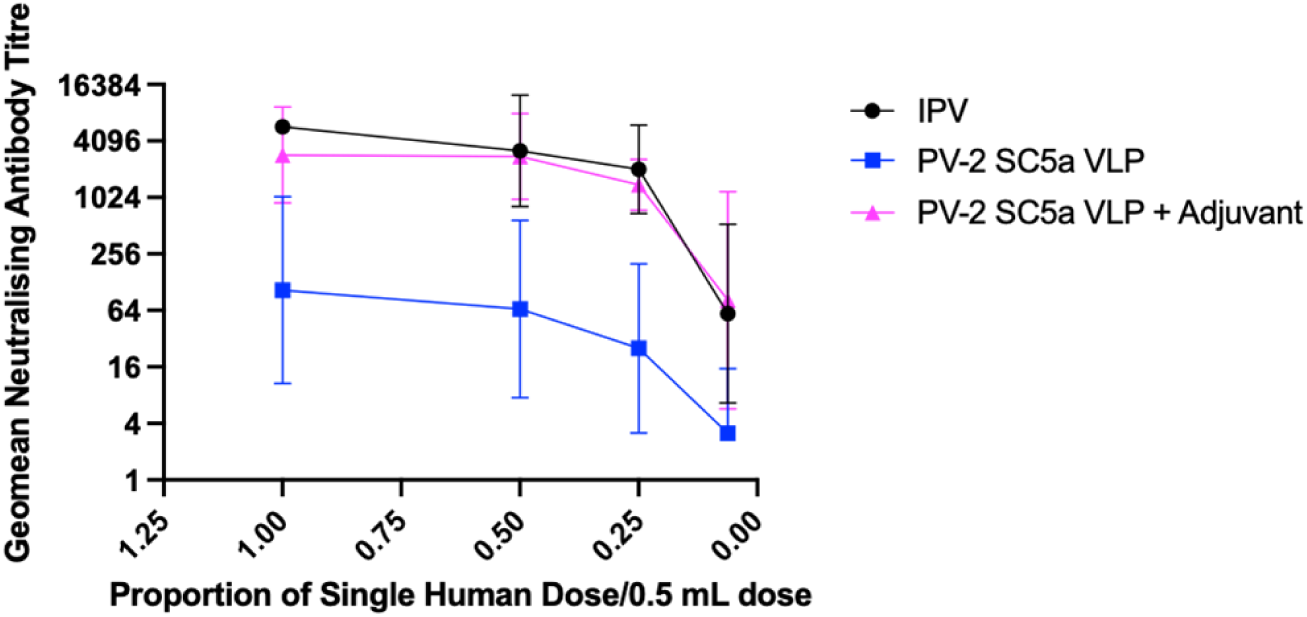
Immunogenicity of PV-2 SC5a VLPs. Dose response in neutralizing antibodies following a single immunization of Wistar rats with PV-2 SC5a VLPs in either the absence or presence of adjuvant. Groups of 10 rats received PV-2 SC5a VLPs at various multiples of human doses and compared to a group of 10 that were immunized with IPV as positive control. Sera were collected 21 dpi and neutralisation titres against the Sabin strain of PV-2 was determined.

As expected, immunisation with IPV produced high levels of neutralising antibody with the titre only falling below 1024 at 0.125 human dose. In the absence of adjuvant, 1 human dose of PV-2 SC5a produced titres of 1024 which reduced as the immunisation dose was lowered. However, in the presence of adjuvant, the response to PV-2 SC5a VLPs equalled the neutralising antibody titres elicited by IPV to 0.25 of a human dose, as observed previously for PV-1 SC6b and PV-3 SC8^38^. Therefore, these data suggest that in the presence of a licensed adjuvant, PV-2 SC5a VLPs are at least as immunogenic as IPV per D Ag.

### Trivalent immunisation induces neutralising antibodies for all three serotypes

The current IPV is administered as a trivalent inoculation, containing inactivated virus particles of all three PV serotypes. It was therefore important to determine if our bioreactor-derived VLPs produced a comparable response when administered as a trivalent mixture.

Rats were immunised with either IPV or a mixture of PV-1 SC6b, PV-2 SC5a and PV-3 SC8 VLPs with or without adjuvant (Alhydrogel 2%) at doses ranging from 1 human dose (32:8:28 D AgU for PV-1, PV-2 and PV-3 respectively) to 0.0625 human dose (2:0.5:1.75 D AgU) and the resulting sera were assessed for neutralising antibody titres (Fig. 6). The trivalent mixture without adjuvant was inferior to IPV for all three serotypes, reflecting our previous observations with monotypic immunisation^38^. However, with adjuvant the immunogenicity was comparable to IPV across all three serotypes. Comparable antibody titres were observed across the dose range, to 0.25 of a single human dose, at which bioreactor-derived VLPs induced antibody titres at levels similar to immunisation with a full human dose. Therefore, this data suggests that in the presence of adjuvant, trivalent immunisation produces comparable immunogenicity to IPV and may offer the potential for dose-sparing.

**Figure 6:**
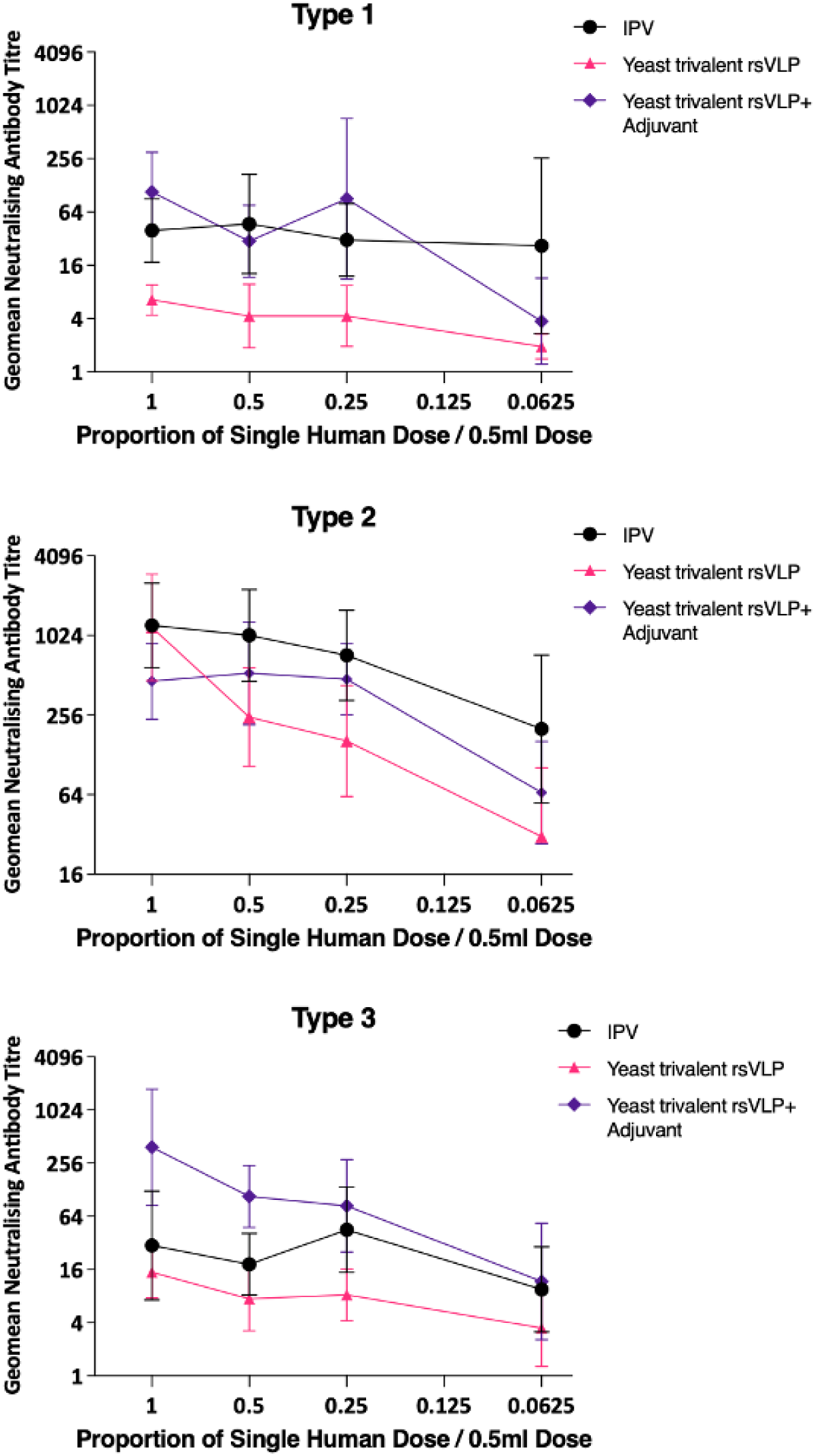
Trivalent immunogenicity of PV VLPs. Dose response in neutralizing antibodies following a single immunization of Wistar rats with trivalent PV VLPs containing PV-1 SC6b, PV-2 SC5a and PV-3 SC8 VLPs, in either the absence or presence of adjuvant. Groups of 10 rats received VLPs at various multiples of human doses and were compared to a group that had received IPV as a positive control. Sera were collected 21 dpi and neutralisation titres against the Sabin strains of PV-1, PV-2 or PV-3 were determined.

## Discussion

While the current PV vaccines, IPV and OPV, have achieved great success, the continued use of OPV has facilitated continued appearance of cVDPVs, which now outnumber wt PV cases year on year. With waning vaccination rates in developed countries sporadic outbreaks of type 2 VDPV have been detected recently in both the UK and the USA^8,9,43^. Improved vaccines are integral to the eradication of PV, illustrated by the success of the recently licenced nOPV2, designed to minimise both reversion to virulence and recombination with other circulating enteroviruses^44^. This, coupled with advances in producing nOPV1 and nOPV3 vaccines, has led to renewed optimism for the eradication of PV^45^. However, given the possibilities of viral escape from vaccine manufacturing facilities together with reversion to virulence (albeit at low frequency) of live vaccines, the complete elimination of PV will require alternative vaccines which do not involve the culture of infectious virus. VLP vaccines produced in recombinant systems may provide the solution and we and others have shown the production of D Ag PV VLPs in several expression systems^32,34–36,38,39^. Our WHO-funded consortium has shown that these recombinant VLPs are as immunogenic as the current IPV in the batch release rat assay^38^. However, to be useful VLP expression and purification must be scaled-up to levels compatible with commercial production. Here, we assess the suitability of yeast-derived PV VLPs for production by fermentation.

Initially, we investigated VLP production using small-scale (250 mL) bioreactor vessels to assess the impact of media, temperature and feed conditions on D Ag yield (Table 1). Following our previous success using flask-based expression, we tested PV-1 VLP expression under conditions of continuous feed, commonly used in fermentation, or bolus-feed, used primarily in flask-based expression. Whilst little difference was seen using BMGY/BMMY medium, YPD/M in continuous-feed conditions led to a ∼1.8-fold increase in D Ag/100 mL culture, whilst minimal medium was not capable of supporting VLP production. Fortuitously, YPD/M is the simplest and cheapest medium of those tested here, suggesting its suitability to produce low-cost PV VLP vaccines - an advantage for production and distribution in LMICs. A range of temperatures had little impact on PV-2 D Ag yield, whilst PV-3 D Ag yield increased markedly at each increment, although this was offset by a decrease in the D:C Ag ratio (Table 1). As a compromise between increased yield whilst maintaining a positive D:C Ag, we selected 28℃ in YPD/M under continuous feed conditions for 10 L fermentation.

All three serotypes produced good levels of D Ag following larger scale fermentation and the VLPs had typical size and morphology (Fig 2A & 2B). Interestingly, the VLPs produced by fermentation were not contaminated with smaller yeast-derived particles such as, but not limited to, alcohol oxidase (AOX) or fatty acid synthase that we had seen previously using flask-based expression^39,46^. These contaminating particles may have arisen as a result of bolus-fed induction during flask-based expression, particularly in the case of AOX, which is known to be induced in the presence of high concentrations of alcohol. Furthermore, we compared the D Ag yield from flask and bioreactor expression (Fig. 2C). Importantly, the VLP yield for every serotype was in controlled fermentation expression conditions. Since PV-2 SC6b produced lower D Ag yields than PV-1 SC6b or PV-3 SC8 in both flask-based and bioreactor expression conditions (Fig 2C), we investigated expression of another PV-2 thermostable mutant, SC5a^26,41^. PV-2 SC5a produced higher yields of D Ag in both flask and bioreactor expression (Fig. 3C) and was more thermostable than both PV-2 SC6b and IPV (Fig. 3D), making it a more suitable VLP for development as a potential vaccine.

Cryo-EM structures have previously been determined for PV-1 SC6b and PV-3 SC8 VLPs produced in flasks^36,38^. To complement these analyses, we performed single particle cryo-EM of bioreactor produced PV-2 SC6b and PV-2 SC5a VLPs to assess their detailed conformational states. This showed that both VLPs adopt native antigenic (D Ag) conformations, are essentially identical in their overall features to native virus particles and contain a lipid moiety bound in their capsid protomers, consistent in length to the sphingosine modelled in PV-2 SC6b VLPs produced from mammalian and insect cell expression^38^.

The most important pre-requisite of any next-generation PV vaccine is that it elicits the same long-lasting immunity against disease as the current vaccine. Whilst PV-2 SC5a VLPs, were not immunogenic enough to match IPV in the absence of adjuvant, the addition of an adjuvant widely used for human vaccines (Alhydrogel) induced antibody titres equivalent to or better than IPV down to 0.25 human dose (Fig. 5), mirroring the improvement in neutralising antibody titre seen by Hong et al., who investigated the immunogenicity of yeast-derived PV-2 SC5a in mice^41^. These differences in immunogenicity between VLPs and IPV in the absence of adjuvant requires further investigation. It may be a result of the formalin-treatment of IPV boosting the immune response, in comparison to untreated VLPs. It could also be due to the viral genome within IPV particles acting as a pathogen-associated molecular pattern to stimulate the innate immune response leading to an improved humoral response, whereas yeast-derived PV-VLPs do not package measurable amounts of RNA^36^. Furthermore, we assessed the immunogenicity of these VLPs in the context of a trivalent immunisation, reflecting the D Ag content of IPV (Fig. 6). In the presence of adjuvant, the neutralisation titres were equivalent or higher than IPV down to 0.25 human dose, in line with our previous findings following monotypic immunisation with PV-1 SC6b and PV-3 SC8 VLPs^38^ and suggesting that these yeast-derived VLPs have the potential for dose-sparing, further reducing the cost of each vaccine dose produced.

In conclusion, we have demonstrated that yeast-derived VLPs from all three PV serotypes are suitable for production by fermentation using the simplest and most cost-effective media, whilst maintaining antigenic, morphological and the thermal stability characteristics required to be a viable next-generation vaccine candidate for a polio-free world. Furthermore, we have improved the production of stabilised PV-2 VLPs and shown that in the presence of adjuvant, trivalent immunisation with these VLPs may be amenable to dose sparing, which in turn can further drive down the cost of future vaccines. Overall, our data shows that the *Pichia pastoris* recombinant expression system can produce PV VLPs using infrastructure already used in LMICs for other vaccines, and therefore is an excellent candidate to produce next-generation PV vaccines globally.

## Methods

### Vector construction

The constructs for PV-1 SC6b, PV-2 SC6b and PV-3 SC8 expression have been described previously^36,38^. The construct for the production of PV-2 SC5a VLPs, which contains different mutations^26^ was produced by the same method. Briefly, the P1 gene for PV2-SC5a, was amplified by polymerase chain reaction (PCR) from a polymerase deleted version of the infectious clone plasmid and cloned into the pPink-HC expression vector multiple cloning site (MCS) using *EcoR*I and *Fse*I (New England Biolabs (NEB)). The 3CD gene was codon-optimised for *P. pastoris* and designed to include a non-cleavable sequence to reduce the potential toxic effects of 3C in a heterologous expression system^47^. The 3CD gene was then cloned into the pPink-HC expression vector MCS using *EcoR*I and *Fse*I (NEB). For the dual promoter constructs, the region from position 1 to position 1285 of 3CD pPink-HC was amplified by polymerase chain reaction (PCR) with primers inserting a *Sac*II restriction site at both the 5’ and 3’ end of the product. The P1 expression plasmid was linearised by *Sac*II (NEB), and the 3CD PCR product was inserted. Clones were screened for directionality, ensuring the promoters for each protein were in the correct orientation. All PCR steps were carried out with Phusion polymerase (NEB) using the manufacturer’s guidelines.

### Yeast transformation and induction

Plasmids were linearized by *Afl*II digestion (NEB) and then transformed into PichiaPink™ Strain one (Invitrogen, USA) by electroporation as per the manufacturer’s guidelines. Transformed yeast cells were plated on *Pichia* adenine dropout (PAD) selection plates and incubated at 28°C until the appearance of white colonies (3-5 days). To screen for high-expression clones, 8 colonies were randomly selected for small-scale (5 mL) expression experiments as described previously^36,40^. Briefly, colonies were cultured in YPD for 48 h at 28°C and shaking at 250 rpm, when cells were pelleted at 1500 × *g* and resuspended in YPM (methanol 0.5% v/v) to induce protein expression followed by culture for a further 48 h. Cultures were fed methanol to 0.5% v/v 24 h post-induction. To determine expression levels of each clone, the samples were analysed by immunoblotting as described below. For flask-based VLP production, a glycerol stock of a high-expressing clone was used to inoculate 5 mL YPD and cultured for 48 h to high density. Subsequently, 2 mL of this starter culture was added to 200 mL YPD in a 2 L baffled flask and cultured at 28°C at 250 rpm for a further 24 h. Cells were pelleted at 1500 × *g* and resuspended in 200 mL YPM (methanol 0.5% v/v) and cultured for a further 48 h. Cultures were fed methanol to 0.5% v/v 24 h post-induction. After 48 h cells were pelleted at 2000 × *g* and resuspended in breaking buffer (50 mM sodium phosphate, 5% glycerol, 1 mM EDTA, pH 7.4) and frozen prior to processing.

### Ambr250 Fermentation of PV VLPs

The *P. pastoris* yeast strains with high expression levels were selected to produce recombinant PV VLPs by high-density fermentation. Each strain was inoculated into 20 mL starter culture of either YPD, BMGY or MD medium and after incubation at 28 °C, 250 rpm for 24 h, approximately 5% of total bioreactor volume (∼8 mL) was transferred to each 250 mL miniature bioreactor, with an initial fill volume of 162 mL of YPD, BMGY, or MD medium. 12 mini bioreactors were incubated in parallel using the Ambr 250 high-throughput system. Batch phase was conducted at 28°C, pH 5.5 and a Dissolved Oxygen (DO) set point of 30% maintained using a DO cascade: stir speed 600-2300 rpm then O_2_ mix up to 40%. Carbon depletion was observed in all vessels between 19 and 20 h post inoculation, indicated by a DO spike, triggering a continuous feed of either a glycerol (BMGY-containing vessels) or dextrose (YPD or MD-containing vessels) feed at 1.7 mL/h to boost biomass. After 8 h, feeding was ceased to allow for induction by methanol. At the beginning of the induction phase, the temperature was reduced to the defined expression temperature, and a metabolic switch was initiated following glycerol or dextrose depletion, by the addition of a methanol bolus to each vessel to a final concentration of 0.5% v/v. Methanol feeding was conducted either by bolus (1 x 7.34 mL/24h) or constant (1.8 mL/h/L respectively) addition from depletion of the initial methanol bolus, identified by a steady decrease in carbon evolution rate (CER). The yeast cultures were collected by centrifugation (8000 x *g*) 48 h after induction and resuspended in breaking buffer (50 mM sodium phosphate, 5% glycerol, 1 mM EDTA, pH 7.4) and frozen prior to processing.

### 10 L Fermentation of PV VLPs

The highly expressing strains were streaked out on PAD plates and single colonies used to inoculate 50 mL of YPD medium in a 250 mL baffled flask. After 48 h incubation at 28°C, 250 rpm, the culture was inoculated into a second stage flask of 200 mL YPD in 1 L baffled flask at OD600 ∼0.2-0.3 and then incubated for a further 22 h. The Sartorius Biostat B-DCU 10L glass bioreactors were then inoculated with 5% starting volume of culture (400 mL), and process conditions were maintained as described above in for Ambr250 HT fermentation. Stir speed was scaled relative to impeller tip speed. The yeast cultures were collected by centrifugation (8000 x *g*) 48 h after induction and resuspended in breaking buffer (50 mM sodium phosphate, 5% glycerol, 1 mM EDTA, pH 7.4) and frozen prior to processing.

### Purification and concentration of PV VLPs

*P. pastoris* cell suspensions were thawed and lysed using a CF-1 cell disruptor at ∼275 MPa chilled to 4°C following the addition of 0.1% Triton-X 100. The resulting lysates were centrifuged at 5,000 rpm for 30 min to remove the larger cell debris, followed by a 10,000 × *g* for 30 min centrifugation to remove further insoluble material. The resulting supernatants were nuclease treated using 25 U/mL DENARASE® (c-LEcta) for 1.5 h at RT with gentle agitation. The supernatants were mixed with PEG 8000 (20% v/v) to a final concentration of 8% (v/v) and incubated at 4°C overnight. The precipitated proteins were pelleted at 5,000 rpm for 30 min and resuspended in PBS. The solutions were clarified again at 5,000 rpm for 30 min and the supernatants collected for a subsequent 10,000 × *g* centrifugation for 30 min to remove insoluble material. The clarified supernatants were collected and pelleted through 30% (w/v) sucrose cushions at 151,000 × *g* (using a Beckman SW 32 Ti rotor) for 3.5 h at 10°C. The resulting pellets were resuspended in PBS + NP-40 (1% v/v) + sodium deoxycholate (0.5% v/v) and clarified by centrifugation at 10,000 x *g* for x min. The supernatants were collected and purified through 15-45% (w/v) sucrose density gradients by ultracentrifugation at 151,000 x *g* (using 17 mL tubes in a Beckman SW32.1 Ti rotor) for 3 h at 10°C^21^. Gradients were collected in 1 mL fractions from top to bottom and analysed for the presence of VLPs through immunoblotting and ELISA.

Peak gradient fractions as determined by ELISA were concentrated to ∼100 uL in PBS + 20 mM EDTA using 0.5 mL 100 kDa centrifugal concentration filters (Amicon) as per the manufacturer’s instructions.

### Enzyme-linked immunosorbent assay (ELISA)

A sandwich ELISA was used to determine the PV D and C Ag contents of sucrose gradient fractions^48^. Briefly, two-fold dilutions of antigen were captured using a PV serotype-specific polyclonal antibody, and detected using serotype-specific monoclonal antibodies, for D Ag (Mab 234, 1050, and 520 for PV-1, PV-2 and PV-3, respectively) or C Ag (Mab 1588 and 517.3 for PV-1 and PV-3, respectively), followed by anti-mouse peroxidase conjugate^49,50^. BRP (Sigma) was used as the standard for D Ag content in each ELISA. All ELISAs were analysed through Biotek PowerWave XS2 plate reader.

### Thermostability assays

Thermostability of PV rsVLPs was assessed as in previous studies^26^. Briefly, the samples were diluted in DPBS to twice the concentration required to obtain an OD of 1.0 in the D Ag ELISA. Duplicate samples were heated for 10 min at a range of temperatures from 30–60°C then diluted 1:1 with 4% dried milk in DPBS and cooled on ice. D Ag and C Ag content was measured by ELISA. The temperature at which the change from D Ag to C Ag occurred is recorded at the point where native antigenicity is reduced by 50%. Thermostability of the VLPs was assessed by measuring loss of D Ag in ELISA using the PV-2 specific Mab 1050 (NIBSC Product No. 1050).

### Immunogenicity in Rats

Immunogenicity of rsVLP preparations was assessed using pharmacopeial methods established at MHRA (NIBSC) for the release of IPV lots. D Ag content was measured by ELISA and immunogenicity was assessed in female Wistar rats^42^. Groups of 10 rats per dose were immunised i.m. with 0.25 ml in each hind leg and terminal bleeds collected on day 21. Sera were analysed for neutralising antibody responses. The neutralising antibody responses to a range of antigen doses were compared to those elicited by a concurrently tested International Standard preparation.

Immunogenicity of rsVLP preparations formulated with adjuvant was assessed in the same way. Prior to inoculation of rats, samples were mixed with 1/10^th^ volume of Alhydrogel (2%, InvivoGen) and agitated for 30 min, 100% of the D Ag was adsorbed onto the aluminium hydroxide.

### Negative stain electron microscopy

To prepare samples for negative stain transmission EM, carbon-coated 300-mesh copper grids were glow-discharged in air at 10 mA for 30 seconds. 3 μl aliquots of purified VLP stocks were applied to the grids for 30 seconds, then excess liquid was removed by blotting. Grids were washed twice with 10 μl distilled H_2_O. Grids were stained with 10 μl 1% uranyl acetate solution, which was promptly removed by blotting before another application of 10 μl 1% uranyl acetate solution for 30 seconds. Grids were subsequently blotted to leave a thin film of stain, then air-dried. EM was performed using an FEI Tecnai G2-Spirit transmission electron microscope (operating at 120 kV with a field emission gun) with a Gatan Ultra Scan 4000 CCD camera (ABSL, University of Leeds).

### Negative stain image processing

Raw micrographs were visualised with ImageJ 1.51d^51,52^.

### Cryo-EM sample preparation and data collection

Sucrose gradient purified fractions of bioreactor-derived PV-2 SC6b and PV-2 SC5a were pooled and concentrated with buffer exchange into PBS + 20mM EDTA (pH 7) using Amicon Ultra centrifugal filter devices (100 kDa MWCO, Merck Millipore) to a final concentration of 0.1 mg/ml (PV-2 SC6b) and 0.6 mg/ml (PV-2 SC5a). For PV-2 SC6b and PV-2 SC5a three microliters of sample were applied to either glow-discharged Quantifoil R2/1 holey carbon support film grids (product No. AGS174-1, Agar Scientific), or Quantifoil R2/1 with 2 nm continuous carbon grids (product No. AGS174-1-2CL, Agar Scientific), respectively. Samples were incubated on the grid for between 30-60 s followed by mechanical blotting and rapid vitrification in a liquid nitrogen cooled ethane/propane slurry with a Vitrobot Mark IV plunge-freezing device (Thermo Fisher Scientific) operated at 4°C and ∼100% relative humidity using a blot force of −15 and blot time of 3.5 s.

For PV-2 SC6b cryo-EM data were acquired at 300 kV with a Titan Krios microscope (Thermo Fisher Scientific) equipped with a Gatan K3 direct electron detector (DED) and a GIF Quantum energy filter (Gatan) operating in zero-loss mode (20 eV slit width), at the electron Bio-Imaging Centre (eBIC), Diamond Light Source, UK. Micrographs were collected as movies using a defocus range of −2.3 μm to −0.8 μm in single-electron counting mode at a nominal magnification of ×105,000 and a calibrated pixel size of 0.831 Å per pixel. For PV-2 SC5a data were collected at 300 kV on a Titan Krios microscope equipped with a K3 DED and a GIF Quantum energy filter mode (20 eV slit width), at The Central Oxford Structural Molecular Imaging Centre (COSMIC), UK. Movies were collected at a nominal magnification of ×105,000 using a defocus range of −2.3 μm to −0.5 μm. The K3 DED was operated in super-resolution mode with a pixel size of 0.830 Å per pixel (0.415 Å per super-resolution pixel). All data were collected using ThermoFisher EPU software, and acquisition parameters are summarized in Table S2.

### Cryo-EM image processing

Data processing and single-particle reconstruction for both the PV-2 SC6b and PV-2 SC5a VLPs was performed using CryoSparc v4.5.3^53^, following standard procedures for icosahedral reconstruction. Raw movies were aligned with Patch Motion Correction and CTF parameters estimated using Patch-CTF. Poor quality images exhibiting significant drift, astigmatism or ice rings were discarded using the manual curation tool. Particles were initially blob picked from a subset of images and subjected to a first 2D classification job to generate suitable templates, which were subsequently used to complete particle picking on the whole data set. Two-dimensional classification was performed iteratively at least twice to clean out junk particles, followed by the generation of five *ab initio* models with the application of icosahedral symmetry. Heterogeneous refinement with icosahedral symmetry was used to further refine the best aligned particle sets to a single good-looking class. These particles were then subjected to homogeneous refinement with icosahedral symmetry and combined with CTF refinement and higher order aberration correction. Final resolution was estimated using the gold-standard FSC 0.143 cut-off on maps output after automatic sharpening and local resolution estimation. Data processing statistics are summarized in Table S1.

### Atomic model building, refinement and analysis

For both PV-2 SC6b and PV-2 SC5a reconstructions the atomic coordinates of the X-ray structure of PV2 (PDB 1EAH) were manually placed into the cryo-EM electron potential maps using UCSF Chimera^54^. Manual fitting was optimised with the UCSF Chimera ‘Fit in Map’ command^54,55^ and the ‘Rigid Body Fit Molecule’ function in Coot^55^. For all structures the cryo-EM map covering six neighbouring capsid protomers (each composed of subunits VP0, VP1 and VP3) was extracted using phenix.map_box within Phenix^56^. Manual rebuilding was performed on the central protomer model using the tools in Coot^55^ and non-crystallographic symmetry operators were used to generate neighbouring protomers, followed by iterative positional and B-factor refinement in real-space using phenix.real_space_refine^57^ within Phenix^56^ to ensure stable refinement of protomer interfaces and minimisation of clashes. All refinement steps were performed in the presence of hydrogen atoms. Only atomic coordinates were refined; the maps were kept constant and each round of model optimization was guided by cross-correlation between the map and the model. Final models were validated using MolProbity^58^, EMRinger^59^ and CaBLAM^60^ integrated within Phenix^56^. Refinement statistics are shown in Table S2.

Structural superpositions and RMSD calculations were performed using the ‘LSQ superpose’ and ‘SSM superpose’ tools within Coot^61^. Molecular graphics were rendered using UCSF ChimeraX^62^.

## Authors and Contributions

LS, MWB, SW, AJM, DJR & NJS conceived and designed the experiments. LS, MWB, KG, JJS, LQ, HF, SM, SC, SW, & NC conducted the experiments. LS, MWB, KG, JJS, HF, SC, SW, SR, NC, EEF, DIS, AJM, DJR & NJS analysed the data. LS, MWB, DJR & NJS wrote the initial manuscript. KG, JJS, NC, SR, CP, JL, AJM, EEF & DIS reviewed and edited the manuscript. Funding was secured for this research by LS, EEF, DIS, AJM, DJR and NJS.

## Data deposition

The atomic coordinates for the cryo-EM structures in this study have been submitted to the Protein Data Bank under the following accession codes (PDB ID): PV-2 SC6b (9H93) and PV-2 SC5a (9H94). The cryo-EM electron potential maps have been deposited in the Electron Microscopy Data Bank under the following accession codes (EMD ID): PV-2 SC6b (EMD-51951) and PV-2 SC5a (EMD-51952).

## Acknowledgments

We thank other members of the Stonehouse/Rowlands group at the University of Leeds for their insightful contributions. This work was performed as part of a WHO-funded collaborative project towards the production of a virus-free polio vaccine involving the following institutions: University of Leeds, University of Oxford, University of Reading, University of Florida, Harvard University, John Innes Centre, the Pirbright Institute and the National Institute for Biological Standards and Control (now the Medicines and Healthcare products Regulatory Agency –MHRA). DIS and EEF are also supported by the UKRI MRC (MR/N00065X/1). We acknowledge Vax-Hub for their collaborative and financial support (EP/R013764/1).

Computation used the Oxford Biomedical Research Computing (BMRC) facility, a joint development between the Wellcome Centre for Human Genetics and the Big Data Institute supported by Health Data Research UK and the National Institute for Health (NIHR) Oxford Biomedical Research Centre. Financial support was provided by the Wellcome Trust Core Award Grant Number 203141/Z/16/Z. We acknowledge Diamond for access and support of the cryo-EM facilities at the UK national electron Bio-Imaging Centre (eBIC), proposal BI28713_11. We thank Dr Rishi Matadeen at the Central Oxford Structural Molecular Imaging Centre (COSMIC), Biochemistry Department, University of Oxford. COSMIC is supported by the Wellcome Trust (#201536), The EPA Cephalosporin Trust, The Wolfson Foundation and a Royal Society/Wolfson Foundation Laboratory Refurbishment Grant (#WL160052). We are grateful for technical assistance from the eBIC and COSMIC staff.

## Ethics declaration

All animal experiments were performed under licenses granted by the UK Home Office under the Animal (Scientific Procedures) Act 1986 revised 2013 and reviewed by the internal NIBSC Animal Welfare and Ethics Review Board. The rat immunogenicity experiments were performed under Home Office licences PPL 70/8979, PPL 80/2478, PPL 80/2050, PPL 80/2537, P30D4C513, PP6108158, P856F6831 and P4F343A03.

## Competing interests

AJM declares he is the named inventor on International Patent Application No. PCT/GB2018/050129 (WO2018134584A1) and granted patent US20190358315A1 which cover the mutations used to stabilise the VLPs.

## Funding

This work was funded via: WHO 2019/883397-O “Generation of virus free polio vaccine – phase IV”.

EPSRC: EP/R013764/1 – “Investigating scalability of a next-generation vaccine for poliomyelitis”.

Vax-Hub Feasibility Study – Scale-up of bioproduction process for a novel virus-like particle (VLP) vaccine candidate for poliomyelitis.

## Notes

### Competing Interest Statement

The authors have declared no competing interest.

## References

1. Polio Now – GPEI. http://polioeradication.org/polio-today/polio-now/.

2. Zambon, M. & Martin, J. Polio eradication: next steps and future challenges. Eurosurveillance 23, (2018).

3. Two out of three wild poliovirus strains eradicated. https://www.who.int/news-room/feature-stories/detail/two-out-of-three-wild-poliovirus-strains-eradicated.

4. Bandyopadhyay, A. S., Garon, J., Seib, K. & Orenstein, W. A. Polio vaccination: past, present and future. Future Microbiol 10, 791–808 (2015).

5. Duizer, E., Ruijs, W. L., van der Weijden, C. P. & Timen, A. Response to a wild poliovirus type 2 (WPV2)-shedding event following accidental exposure to WPV2, the Netherlands, April 2017. Eurosurveillance 22, 30542 (2017).

6. EU Threats.

7. Minor, P. Vaccine-derived poliovirus (VDPV): Impact on poliomyelitis eradication. Vaccine 27, 2649–2652 (2009).

8. Klapsa, D. et al. Sustained detection of type 2 poliovirus in London sewage between February and July, 2022, by enhanced environmental surveillance. The Lancet 400, 1531–1538 (2022).

9. Tanne, J. H. Polio emergency declared in New York State over virus found in wastewater. BMJ 378, o2211 (2022).

10. Jiang, P. et al. Evidence for emergence of diverse polioviruses from C-cluster coxsackie A viruses and implications for global poliovirus eradication. Proc Natl Acad Sci U S A 104, 9457– 9462 (2007).

11. Jegouic, S. et al. Recombination between Polioviruses and Co-Circulating Coxsackie A Viruses: Role in the Emergence of Pathogenic Vaccine-Derived Polioviruses. PLoS Pathog 5, e1000412 (2009).

12. Greene, S. A. et al. Progress Toward Polio Eradication — Worldwide, January 2017–March 2019. MMWR Morb Mortal Wkly Rep 68, 458–462 (2019).

13. Lulla, V. et al. An upstream protein-coding region in enteroviruses modulates virus infection in gut epithelial cells. Nat Microbiol 4, 280–292 (2019).

14. Tuthill, T. J., Groppelli, E., Hogle, J. M. & Rowlands, D. J. Picornaviruses. in Current topics in microbiology and immunology vol. 343 43–89 (2010).

15. Molla, A. et al. Stimulation of poliovirus proteinase 3Cpro-related proteolysis by the genome-linked protein VPg and its precursor 3AB. J Biol Chem 269, 27015–20 (1994).

16. Jore, J., De Geus, B., Jackson, R. J., Pouwels, P. H. & Enger-Valk, B. E. Poliovirus Protein 3CD Is the Active Protease for Processing of the Precursor Protein P1 in vitro. Journal of General Virology 69, 1627–1636 (1988).

17. Kräusslich, H. G., Nicklin, M. J., Lee, C. K. & Wimmer, E. Polyprotein processing in picornavirus replication. Biochimie 70, 119–30 (1988).

18. Jiang, P., Liu, Y., Ma, H.-C., Paul, A. V & Wimmer, E. Picornavirus morphogenesis. Microbiol Mol Biol Rev 78, 418–37 (2014).

19. Hogle, J. M., Chow, M. & Filman, D. J. Three-dimensional structure of poliovirus at 2.9 A resolution. Science 229, 1358–65 (1985).

20. Filman, D. J. et al. Structural factors that control conformational transitions and serotype specificity in type 3 poliovirus. EMBO J 8, 1567–1579 (1989).

21. Basavappa, R. et al. Role and mechanism of the maturation cleavage of VP0 in poliovirus assembly: Structure of the empty capsid assembly intermediate at 2.9 Å resolution. Protein Science 3, 1651–1669 (1994).

22. Le Bouvier, G. L. The modification of poliovirus antigens by heat and ultraviolet light. Lancet 269, 1013–6 (1955).

23. Strauss, M., Schotte, L., Karunatilaka, K. S., Filman, D. J. & Hogle, J. M. Cryo-electron Microscopy Structures of Expanded Poliovirus with VHHs Sample the Conformational Repertoire of the Expanded State. J Virol 91, (2017).

24. Le Bouvier, G. L. Poliovirus D and C antigens: their differentiation and measurement by precipitation in agar. Br J Exp Pathol 40, 452–463 (1959).

25. Beale, A. J. & Mason, P. J. The measurement of the D-antigen in poliovirus preparations. Journal of Hygiene 60, 113–121 (1962).

26. Fox, H., Knowlson, S., Minor, P. D. & Macadam, A. J. Genetically Thermo-Stabilised, Immunogenic Poliovirus Empty Capsids; a Strategy for Non-replicating Vaccines. PLoS Pathog 13, e1006117 (2017).

27. McAleer, W. J. et al. Human hepatitis B vaccine from recombinant yeast. Nature 307, 178–80 (1984).

28. Sasagawa, T. et al. Synthesis and assembly of virus-like particles of human papillomaviruses type 6 and type 16 in fission yeast Schizosaccharomyces pombe. Virology 206, 126–35 (1995).

29. Smith, J., Lipsitch, M. & Almond, J. W. Vaccine production, distribution, access, and uptake. Lancet 378, 428–38 (2011).

30. Ansardi, D. C., Porter, D. C. & Morrow, C. D. Coinfection with recombinant vaccinia viruses expressing poliovirus P1 and P3 proteins results in polyprotein processing and formation of empty capsid structures. J Virol 65, 2088–92 (1991).

31. Viktorova, E. G. et al. Newcastle Disease Virus-Based Vectored Vaccine against Poliomyelitis. J Virol 92, (2018).

32. Bahar, M. W. et al. Mammalian expression of virus-like particles as a proof of principle for next generation polio vaccines. NPJ Vaccines 6, (2021).

33. Bräutigam, S., Snezhkov, E. & Bishop, D. H. L. Formation of Poliovirus-like Particles by Recombinant Baculoviruses Expressing the Individual VP0, VP3, and VP1 Proteins by Comparison to Particles Derived from the Expressed Poliovirus Polyprotein. Virology 192, 512–524 (1993).

34. Xu, Y. et al. Virus-like particle vaccines for poliovirus types 1, 2, and 3 with enhanced thermostability expressed in insect cells. Vaccine 37, 2340–2347 (2019).

35. Marsian, J. et al. Plant-made polio type 3 stabilized VLPs—a candidate synthetic polio vaccine. Nat Commun 8, 245 (2017).

36. Sherry, L. et al. Production and Characterisation of Stabilised PV-3 Virus-like Particles Using Pichia pastoris. Viruses 14, 2159 (2022).

37. Kim, H. J. & Kim, H.-J. Yeast as an expression system for producing virus-like particles: what factors do we need to consider? Lett Appl Microbiol 64, 111–123 (2017).

38. Sherry, L. et al. Risk-free polio vaccine: Recombinant expression systems for production of stabilised virus-like particles 2 3.

39. Sherry, L. et al. Protease-Independent Production of Poliovirus Virus-like Particles in Pichia pastoris: Implications for Efficient Vaccine Development and Insights into Capsid Assembly. Microbiol Spectr (2022) doi:10.1128/spectrum.04300-22.

40. Sherry, L. et al. Comparative Molecular Biology Approaches for the Production of Poliovirus Virus-Like Particles Using Pichia pastoris. mSphere 5, (2020).

41. Hong, Q. et al. Vaccine Potency and Structure of Yeast-Produced Polio Type 2 Stabilized Virus-like Particles. Vaccines (Basel) 12, 1077 (2024).

42. Van Steenis, G., Van Wezel, A. L. & Sekhuis, V. M. Potency testing of killed polio vaccine in rats. Dev Biol Stand **Vol.** 47, 119–128 (1980).

43. Iacobucci, G. UK childhood vaccination rates fell last year in almost all programmes, figures show. BMJ 378, o2353 (2022).

44. Yeh, M. Te et al. Engineering the Live-Attenuated Polio Vaccine to Prevent Reversion to Virulence. Cell Host Microbe 27, 736–751.e8 (2020).

45. Yeh, M. Te et al. Genetic stabilization of attenuated oral vaccines against poliovirus types 1 and 3. Nature 619, 135 (2023).

46. Snowden, J. S. et al. Structural insight into Pichia pastoris fatty acid synthase. Sci Rep 11, (2021).

47. Barco, A., Feduchi, E. & Carrasco, L. Poliovirus Protease 3Cpro Kills Cells by Apoptosis. Virology 266, 352–360 (2000).

48. Singer, C. et al. Quantitation of poliovirus antigens in inactivated viral vaccines by enzyme-linked immunosorbent assay using animal sera and monoclonal antibodies. J Biol Stand 17, 137–50 (1989).

49. Minor, P. D. et al. Antigenic structure of chimeras of type 1 and type 3 polioviruses involving antigenic sites 2, 3 and 4. Journal of General Virology 72, 2475–2481 (1991).

50. Ferguson, M., Wood, D. J. & Minor, P. D. Antigenic structure of poliovirus in inactivated vaccines. Journal of General Virology 74, 685–690 (1993).

51. Schindelin, J., et al. Fiji: an open-source platform for biological-image analysis. Nat Methods 9, 676–682 (2012).

52. Schneider, C. A., Rasband, W. S. & Eliceiri, K. W. NIH Image to ImageJ: 25 years of image analysis. Nat Methods 9, 671–5 (2012).

53. Punjani, A., Rubinstein, J. L., Fleet, D. J. & Brubaker, M. A. CryoSPARC: Algorithms for rapid unsupervised cryo-EM structure determination. Nat Methods 14, 290–296 (2017).

54. Pettersen, E. F. et al. UCSF Chimera - A visualization system for exploratory research and analysis. J Comput Chem 25, 1605–1612 (2004).

55. Casañal, A., Lohkamp, B. & Emsley, P. Current developments in Coot for macromolecular model building of Electron Cryo-microscopy and Crystallographic Data. Protein Science 29, 1069–1078 (2020).

56. Liebschner, D. et al. Macromolecular structure determination using X-rays, neutrons and electrons: Recent developments in Phenix. Acta Crystallogr D Struct Biol 75, 861–877 (2019).

57. Afonine, P. V. et al. Real-space refinement in PHENIX for cryo-EM and crystallography. Acta Crystallogr D Struct Biol 74, 531–544 (2018).

58. Williams, C. J. et al. MolProbity: More and better reference data for improved all-atom structure validation. Protein Science 27, 293–315 (2018).

59. Barad, B. A. et al. EMRinger: Side chain-directed model and map validation for 3D cryo-electron microscopy. Nat Methods 12, 943–946 (2015).

60. Prisant, M. G., Williams, C. J., Chen, V. B., Richardson, J. S. & Richardson, D. C. New tools in MolProbity validation: CaBLAM for CryoEM backbone, UnDowser to rethink “waters,” and NGL Viewer to recapture online 3D graphics. Protein Science 29, 315–329 (2020).

61. Emsley, P., Lohkamp, B., Scott, W. G. & Cowtan, K. Features and development of Coot. Acta Crystallogr D Biol Crystallogr 66, 486–501 (2010).

62. Pettersen, E. F. et al. UCSF ChimeraX: Structure visualization for researchers, educators, and developers. Protein Science 30, 70–82 (2021).

