## Supplementary Materials for "Upscaling production of immunogenic poliovirus virus-like particles in *Pichia Pastoris* by controlled fermentation"

**Supplementary Tables**

|  | PV-2 SC6b | PV-2 SC5a |
| --- | --- | --- |
|  | EMD-51951 | EMD-51952 |
| **Data Collection** |  |  |
| Microscope | Titan Krios (eBIC) | Titan Krios (COSMIC) |
| Voltage (kV) | 300 | 300 |
| Detector | Gatan K3 | Gatan K3 |
| Recording mode | Counting | Super resolution |
| Nominal magnification (×) | 105000 | 105000 |
| Pixel size (Å) (super-resolution) | 0.831 | 0.830 (0.415) |
| Defocus range (µm) | -2.3 to -0.8 | -2.3 to -0.5 |
| Dose rate (*e^−^*/pixel/s) | 15.16 | 10.80 |
| Frames per movie | 35 | 50 |
| Movie exposure time (s) | 1.6 | 3.0 |
| Total electron dose (*e^−^*/Å^2^) | 35.12 | 47.50 |
| **Data processing** |  |  |
| Movies | 29172 | 1862 |
| Initial particles (no.) | 660747 | 243874 |
| Final particles (no.) | 113853 | 147409 |
| Box size (pixels) | 576 | 576 |
| Symmetry | I1 | I1 |
| Resolution (Å) | 2.4 | 2.1 |
| Map sharpening *B*-factor (Å^2^) | -86.1 | -94.4 |

**Table S1: Structure refinement and validation for the capsid protein (VP0, VP1, VP3)**

|  | PV-2 SC6b | PV-2 SC5a |
| --- | --- | --- |
|  | PDB 9H93 | PDB 9H94 |
| **Model composition** |  |  |
| Non-hydrogen atoms | 5530 | 5902 |
| Protein residues | 703 | 724 |
| Ligands | SPH: 1 | SPH: 1 |
| Waters |  | 198 |
| **Refinement** |  |  |
| Resolution (Å) | 2.4 | 2.1 |
| Map CC^a^ (Mask) | 0.89 | 0.89 |
| Map CC^a^ (Volume) | 0.81 | 0.81 |
| **RMS deviations** |  |  |
| Bond lengths (Å) | 0.004 | 0.003 |
| Bond angles (°) | 0.622 | 0.579 |
| **Mean B-factor (Å^2^)** |  |  |
| Protein | 20.40 | 13.41 |
| Ligand | 20.73 | 17.78 |
| Water |  | 15.22 |
| **Validation** |  |  |
| Molprobity^b^ score (percentile) | 1.29 (98^th^) | 1.18 (99^th^) |
| Clashscore^b^, all atoms (percentile) | 2.56 (98^th^) | 3.10 (98^th^) |
| Ramachandran favoured (%) | 97.11 | 97.95 |
| Ramachandran allowed (%) | 2.56 | 2.05 |
| Ramachandran outliers (%) | 0.00 | 0.00 |
| Rotamer favoured (outliers) (%) | 94.68 (1.33) | 97.27 (0.46) |
| Cβ deviations >0.25 Å (%) | 0.00 | 0.00 |
| CaBLAM outliers (%) | 1.2 | 0.9 |
| CA Geometry outliers (%) | 0.29 | 0.28 |
| **EMRinger^c^ score** | 5.89 | 7.61 |

**Table S2: Structure refinement and validation for the capsid protein (VP0, VP1, VP3)**

^a^Map CC is given for the full particle reconstruction.

^b^Williams *et al.* (2018) Protein Sci 27:293-315.

^c^Barad *et al.* (2015) Nature Methods 12:943–946.

**Supplementary Figures**

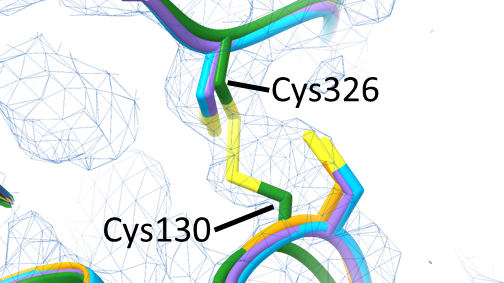

**Figure S1: Bioreactor produced PV-2 SC6b disulphide bond between Cys130 – Cys326 of the capsid protomer subunit VP0.** Close up view of the Cys130 and Cys326 residues in subunit VP0 of PV-2 SC6b (equivalent to Cys61 and Cys257 in the mature VP2 sequence numbering), shown as sticks and coloured green. The electron potential map between Cys130-Cys326 supports the formation of a disulphide bond between the amino acid residues. Map is shown as a wire mesh at a threshold of 1.5 σ. The structures of related PV-2 VLPs are shown superposed in orange (PV-2 SC5a), purple (PV-2 SC6b from mammalian cell expression) and cyan (PV-2 SC6b from insect cell expression) (Ref. 38)
